# Resveratrol prevents amyloid fibrillation of insulin by arresting it in a bioactive oligomeric form

**DOI:** 10.1101/611376

**Authors:** Bani Kumar Pathak, Debajyoti Das, Sayan Bhakta, Partha Chakrabarti, Jayati Sengupta

## Abstract

Insulin fibrillation is a limiting factor for its long-term storage because of considerably reduced bioavailable moieties. Deposition of fibrillated insulin can also cause subcutaneous insulin amyloidoma. Toxic phenolic compounds along with Zinc are used during commercial preparation of insulin to stabilize it in a hexameric form. Designed or repurposed natural small molecules with anti-amyloidogenic properties could thus be attractive agents for preventing insulin fibrillation. Natural polyphenolic compounds which have been shown to serve as anti-amyloid agents for proteins associated with neurodegenerative diseases are potential candidates for such function.

In this study we have demonstrated that resveratrol, a natural polyphenol, can not only prevent insulin fibrillation but can also preserve insulin in a bioactive oligomeric form even at high temperature. While investigating the influence of some natural polyphenols on human insulin (hINS) in a condition inductive to amyloid fibrillation at physiological pH, we found attenuation, to different extents, of insulin fibril formation. However, visualization of polyphenol-treated hINS revealed that resveratrol in particular has the unique ability to arrest hINS before the onset of fibrillation growth in soluble oligomeric forms with discrete spherical morphology. Importantly, insulin treated with resveratrol retains its full biological activity *in vivo* and exerts no toxicity towards cell lines. Structural characterization of the major population of resveratrol-induced insulin oligomers by cryo-EM and single particle 3D reconstruction revealed its morphology that resembled crystal structure of insulin hexamer formulated with phenolic compounds. Thus, our study suggests that resveratrol can be an effective nontoxic substituent of phenolic compounds for insulin preservation.

## Introduction

The anti-hyperglycemic hormone insulin is widely used as a drug for type 1 and in some cases type 2 diabetes. It is a 51-residue globular protein consisting of 21 residues long A-chain and 30 residues long B-chain that are connected by 2 disulfide bridges [1] (Figure S1). The structural analysis confirms that the A-chain contains a helical hairpin whereas B-chain consists of a single helix [2]. In the pancreatic β-cells, insulin is stored as an inactive hexameric assembly stabilized by zinc ions [3, 4] and following secretion into blood, high dilution and change of pH induce its dissociation into dimer and subsequently into a biologically active monomer [5]. In solution, insulin apparently coexists with a dimer, tetramer, hexamer, or other higher-order multimers in equilibrium [6]. Although monomeric insulin is the active form of the hormone, insulin monomer is less stable than the Zn^2+^-coordinated hexameric form and is prone to aggregation [7]. Mechanistically, the unfolded insulin monomer with exposed hydrophobic regions triggers amyloidogenesis [8], with the typical cross-β structure as that of mature fibrils of other disease causing amyloids [9], and thus belongs to the category of natively structured amyloidogenic proteins.

The presence of insulin amyloid fibrils has been reported in patients with diabetes as well as in normal aging [9, 10]. Increasing number of evidence has been accumulated on insulin amyloidoma, a medical condition in diabetes patients where extracellular insulin fibrils are found at the site of insulin injection [11, 12], that may also lead to poor penetration of the injected insulin and variable insulin response [7]. Moreover, insulin fibrillation has been a limiting factor in the long-term storage of insulin, causing reduced bioavailability of the existing insulin formulations [7, 13] and impairment of the efficiency of insulin delivery systems such as insulin pumps [7].

Commercially available insulin analogs are formulated with stabilizing excipients such as m-cresol and other phenolic products to avoid or minimize fibrillation and provide enhanced stability [14, 15]. Nevertheless, recent literature raised a relevant question about the toxicity of insulin formulations. Reports suggest pure m-cresol, phenol, and even insulin formulations, display enhanced cytotoxicity and proinflammatory response, resulting in significant cell death, when they are exposed to certain dosage [16, 17]. Due to their adverse cytotoxic and proinflammatory effects, phenolic derivatives can potentially act as toxins in the kidneys, skin, liver, respiratory, gastrointestinal, cardiovascular and central nervous systems [18]. Several years of exposure to this toxin renders higher susceptibility to develop tumors [19], demonstrating its carcinogenic potential. Thus, repeated dosage of insulin formulations containing m-cresol or other phenolic compounds may raise serious medical concern for diabetic patients. Therefore, any nontoxic molecule that stabilizes insulin oligomers and at the same time permits the release of fast-acting monomeric active form has a potential clinical utility. A very recent study has demonstrated a chemical synthesis approach for producing biologically active insulin variant [20].

Various small molecules, peptides, oligomer- or amyloid-specific antibodies are known to act as effective inhibitors of protein fibrillation [21]. Few studies have reported inhibitory effects of small molecules such as epigallocatechin gallate (EGCG), quercetin or quinones on insulin fibrillation [22]. Among the small molecules, naturally occurring polyphenols have definite advantages since they are non-toxic at moderate concentrations and stable in serum without having an immune response [23]. Interestingly, emerging evidence suggests that ingestion of polyphenol-rich pomace improves insulin sensitivity [24, 25].

Here, we have examined the anti-aggregation potential of naturally occurring polyphenols on the fibrillation of human insulin under close to physiological conditions and explored the degree of efficacy of three naturally occurring polyphenolic compounds (Figure S1a-c) as anti-amyloidogenic factors for insulin (Figure S1d,e). Our study reveals that while all polyphenols are effective in reducing insulin fibrillation to a variable extent, the resveratrol in particular, abundantly found in red wine, chocolate and other dietary products, exhibits the strongest effect. More importantly, resveratrol-stabilized insulin has comparable glucose lowering activity to native insulin, suggesting that resveratrol-stabilized oligomers are capable of efficiently releasing active insulin monomers. Our structural analysis reveals that resveratrol-stabilized insulin adopts a tri-lobed globular shape resembling native insulin hexamer conformation.

Unlike fibrillation involved in neurodegenerative diseases, insulin fibrillation not only complicates the disease but also reduces availability of active insulin molecules. Our results demonstrate that resveratrol inhibits the fibrillation process of insulin at high temperature in such a way that active insulin molecules get stabilized. Thus, resveratrol can be a potential nontoxic stabilizer molecule for preserving insulin.

## Results

### Resveratrol inhibits insulin fibrillation differently compared to other polyphenolic compounds

We have investigated insulin fibrillation under conditions closely resembling a physiological one (pH 7.4) [26], in contrast to the common experimental conditions of insulin fibrillation, such as low pH, extremely high temperature, high concentration of salts, and/or the presence of organic solvent [27, 28]. The anti-aggregation properties of three native polyphenols, resveratrol (trans-3, 4, 5-trihydroxystilbene, RES), epigallocatechin gallate (EGCG) and curcumin (CUR), along with flavanone glycoside-hesperidin (HES), were tested for human insulin aggregation using Thioflavin T (Th-T) fluorescence. Insulin, purchased from SIgma, had no phenolic additives. A significant decrease in fluorescence intensities was seen when polyphenols were incubated for 24 h with insulin in the equimolar ratio (Figure S2a). In contrast, another bioflavonoid, HES (which is not a polyphenol), did not show similar effect indicating specific insulin anti-aggregation role of polyphenols.

This observation prompted us to visualize the polyphenol-induced morphological changes during insulin oligomerization. The effect of polyphenols on insulin fibrillation was examined using atomic force microscopy (AFM). The AFM image revealed that insulin fibrils formed in the absence of polyphenols (Figure 1a). In contrast, discrete spherical oligomers were the predominant species observed in the presence of an equimolar concentration of resveratrol to insulin (Figure 1b), whereas a mixed population of insulin oligomers (much larger in size) was found when insulin was treated with EGCG or curcumin (Figure 1c and 1d).

**Figure 1:**
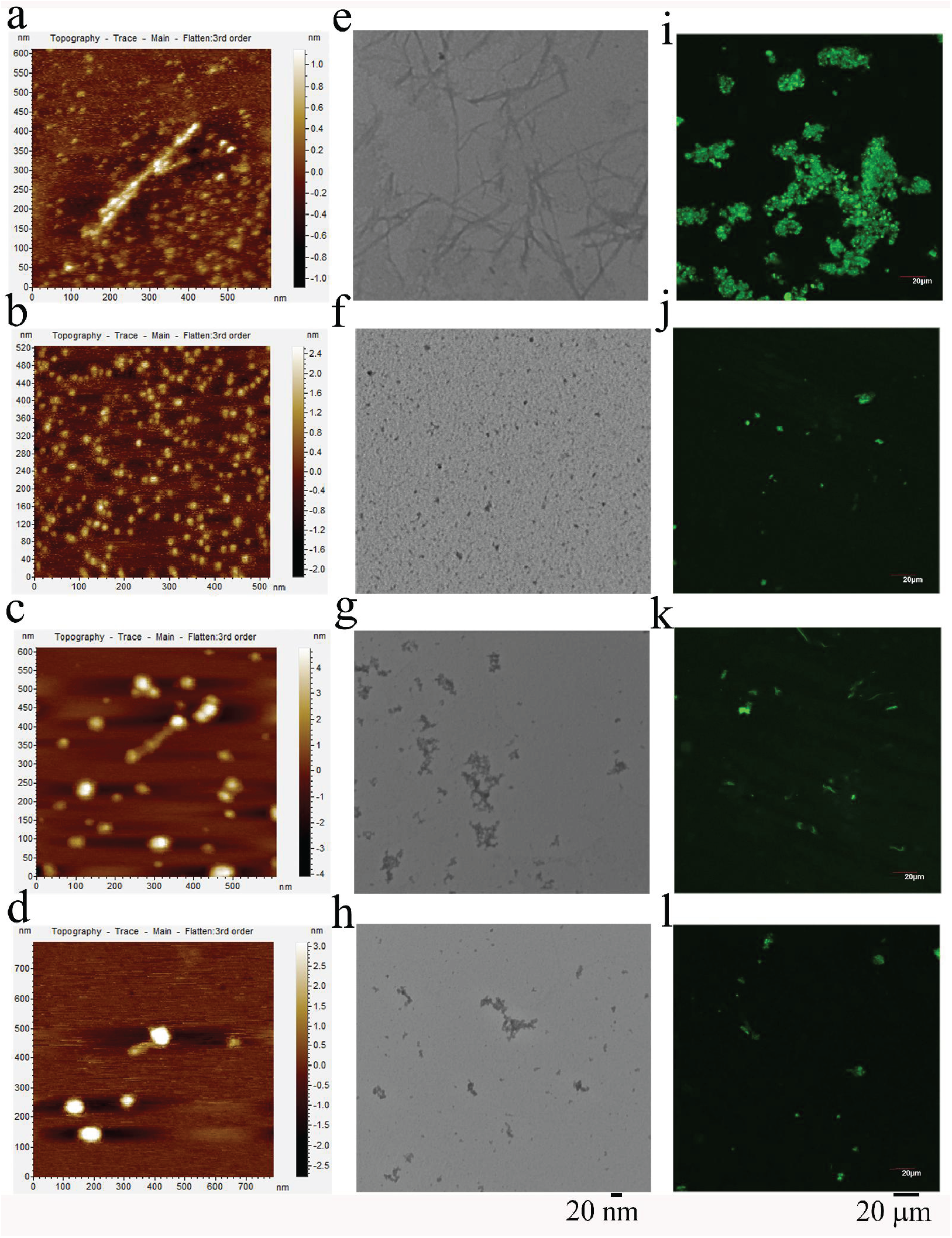
Visualization of insulin fibrillation in the presence and absence of polyphenols. Time-lapse AFM images of insulin fibrillation (a) in the absence, and presence of 100μM of (b) RES, (c) EGCG and (d) CUR are shown. TEM images of insulin in (e) absence and presence of 100μM of (f) RES, (g) EGCG and (h) CUR. AFM and TEM visualizations show that resveatrol stabilized insulin in a homogenious population of small oligomers. Confocal images of insulin fibrillation after 24 hours of incubation (i) in the absence, and presence of 100μM of (j) RES, (k) EGCG and (l) CUR suggest there is less fibril formation in resveratrol-stabilized insulin.

To gain insight into the sizes of insulin oligomers in the presence of polyphenols more precisely, transmission electronic microscopy (TEM) visualization was done. TEM image manifested a mesh of insulin aggregates characterized as linear and complexly intertwined fibrils (Figure 1e). Samples co-incubated with the polyphenolic compounds showed strong anti-aggregation effect (Figure 1f-1h). Interestingly, tiny dotted structure (<20nm), with regular shape and size, was observed only in the presence of resveratrol (Figure 1f). In the presence of EGCG or curcumin, on the other hand, short irregular protofibril-like structures were found buried in the amorphous aggregates (Figure 1g and 1h).

We used Th-T fluorescence to visualize insulin fibrillation under a confocal microscope as well, which also showed significantly lower population of amyloid structures upon polyphenol treatments (Figure 1i-l), particularly in the presence of resveratrol (Figure 1j). It should be noted here that, although imaging using imaging using Th-T fluorescence can detect oligomers with cross β-sheet structures characteristic of amyloid fibers, it lacks the proper resolution to differentiate between fibrillar species present in the aggregates.

Visualizations of polyphenol-mediated anti-fibrillation phenomena revealed remarkable differences in size and shape of insulin aggregates in the presence of resveratrol as compared to other two polyphenols. It seemed from a dose-dependent study that resveratrol is effective when used at 1:0.5 or more of insulin:resveratrol ratio (Figure S3).

### Long-term effect of insulin fibrillation inhibition by the polyphenols

We further sought to investigate long-term effects of the polyphenols on insulin fibrillation. The increase in Th-T fluorescence intensity was observed after 4h of agitation at 65°C and reached the maximum insulin fibrillation at 12h in the absence (Control) of polyphenols (Figure 2a). Polyphenolic compounds, when co-incubated with the insulin, reduced the total yield of fibrils formed after 24h of incubation (Figure 2a). Furthermore, each of the three compounds can increase the amount of unreacted insulin in soluble fraction for up to 70 to 75% of the total protein used in the experiment (Figure 2b). Here, unreacted insulin signifies the amount of insulin retained in soluble fraction after 24h of incubation in the absence or presence of polyphenols.

**Figure 2:**
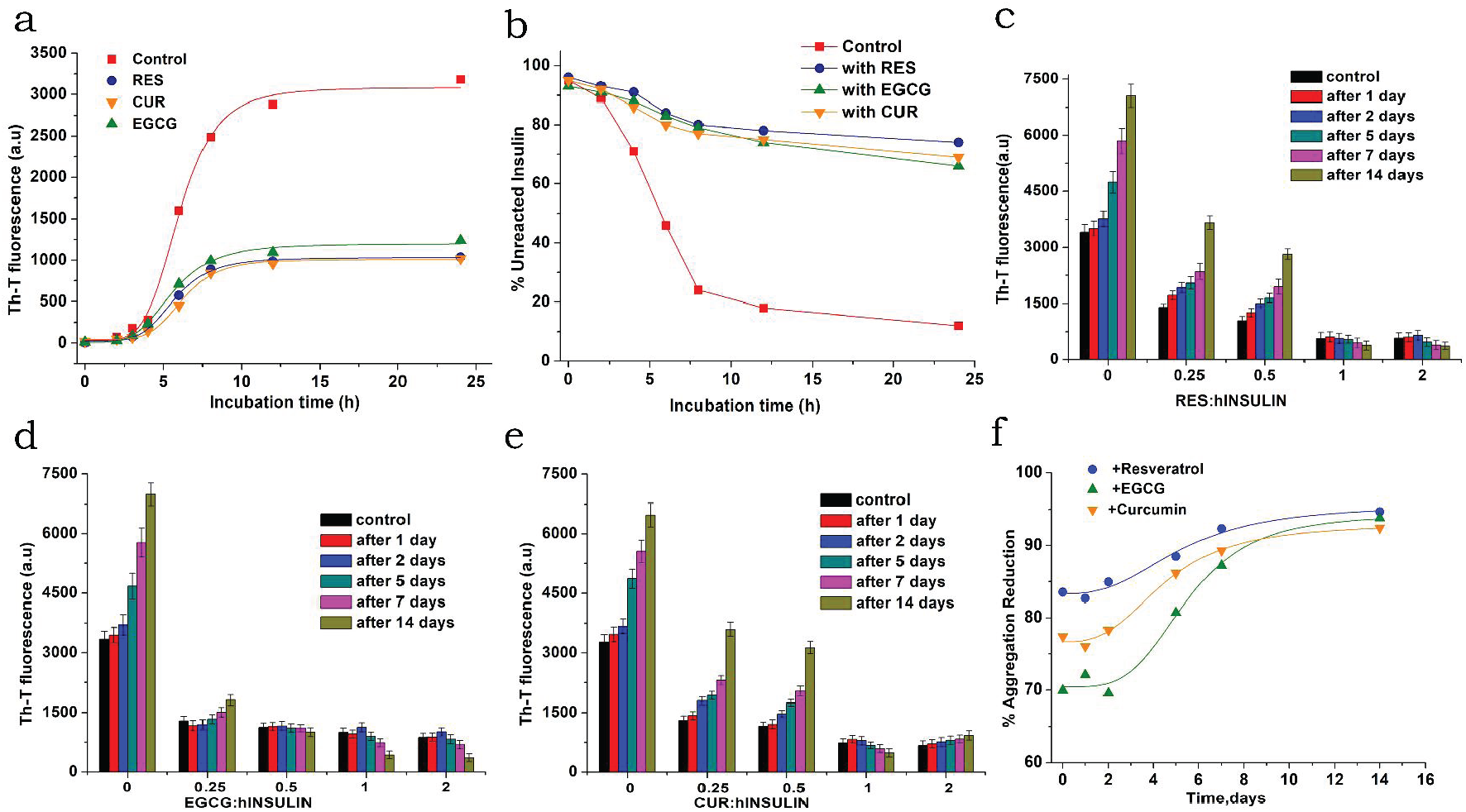
Long-term effects of polyphenols on insulin aggregation. (a) Time-course of insulin fibrillation in absence (control) and presence of three polyphenols, resveratrol (RES), curcumin (CUR) and epigallocatechin gallate (EGCG) as indicated in the diagram by separate symbol and color code. (b) The amount of unreacted insulin was determined with or without (control) RES, EGCG and CUR by Bradford protein assay. (c-e) Insulin fibrilation was monitored by measuring Th-T fluorescence at 485 nm in the absence and in the presence of 25 μM, 50 μM, 50 μM, 100 μM and 200 μM of (c) RES, (d) EGCG and (e) CUR, for upto 14 days at 37°C in PBS after 24h of incubation at 65°C. Separate control experiment (without polyphenols) was performed to study the effect of each phenolic compound on insulin fibrillation for 14 days. Averages of three experiments including three technical repeats are shown (n = 3), with error bars. (f) Polyphenols mediated insulin aggregation reduction was calculated in percent at insulin/polyphenols =1:1 after 24h of incubation. The calculation for percentage insulin aggregation reduction by the polyphenols was performed assuming the Th-T fluorescence value of insulin aggregates without polyphenols to be 100%. Time course experiments were repeated thrice and their average values were taken for final data plotting.

To investigate the long-term effects of the polyphenolic compounds on insulin oligomerization, native insulin (control) and polyphenol-treated (reaction sets in the presence of each of the polyphenolic compounds) samples were incubated at 65°C for 24h and then kept at 37°C for 14 days with Thioflavin T (Th-T) fluorescence measured periodically at different time points (Figure 2c-2e). In the absence of polyphenols, almost double Th-T fluorescence was seen at day 14 (compared to control) indicating a continuation of insulin fibrillation process even at 37°C. Ratios of polyphenols to insluin of more than 0.5:1 showed most promising inhibitory effects on insulin fibrillation event. At equimolar ratio, from the very first day of incubation, resveratrol is much more effective in insulin aggregation reduction compared to other two polyphenols (Figure 2f).

### Resveratrol is non-toxic and effectively allows the release of insulin in its active form

Resveratrol-mediated formation of homogeneous spherical species of insulin prompted us to investigate its biological activity. Insulin fibrillation assay in the presence of resveratrol showed comparable insulin signaling with native insulin as a readout of AKT phosphorylation in HepG2 cells (Figure 3a). Though EGCG prevented insulin fibrillation, it failed to retain the biological activity of insulin, resulting in very little or no induction of AKT phosphorylation.

**Figure 3:**
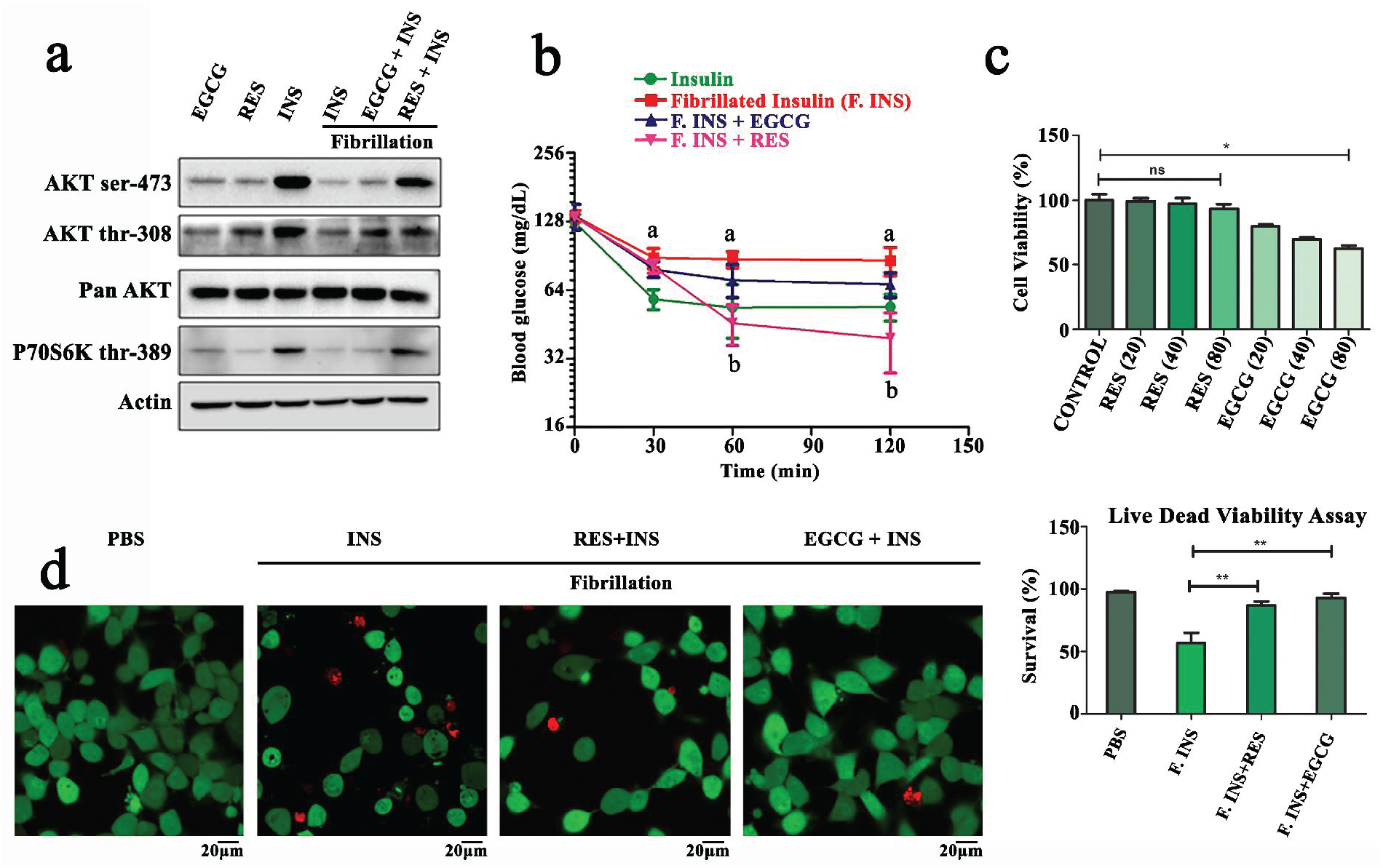
Insulin signaling and cytotoxicity assay. (a) Fibrillated insulin in the absence and presence of RES and EGCG was used to treat HepG2 cells. Insulin signaling assay was based on the Akt phosphorylation along with expression of different control candidate proteins as evident in the given western blotting picture. (b) Insulin Tolerance Test performed in BALB/c mice in four different groups: Insulin, fibrillated insulin, fibrillated insulin with EGCG and fibrillated insulin with resveratrol (5 mice per group). a = statistically significant difference in blood glucose level between the native insulin-treated and fibrillated insulin-treated group, b = statistically significant difference in blood glucose level between fibrillated insulin and fibrillated insulin with resveratrol-treated group. (c) MTT Assay was performed to determine cytotoxicity. Cells were treated with resveratrol and EGCG with varying concentration of 20, 40 and 80 μM for 24 hrs. (d) Cell viability was further assessed using the Live Dead viability assay. Red colored cells are dead ones, green colored are live. Statistical analysis was performed to determine the percent survival rate of HepG2 cells upon treatment. Data are represented as mean±SD; *, p<0.05.

Further, to validate the more direct physiological role of insulin, we performed the ‘Insulin Tolerance Test’ (ITT) in BALB/c mice. Resveratrol-treated insulin showed a comparable lowering of blood glucose with native insulin *in vivo*, whereas fibrillated insulin and EGCG treated insulin showed very little change in blood glucose level (Figure 3b). Moreover, resveratrol-mediated stabilization of insulin followed different kinetics as revealed by delayed onset of glucose-lowering response in comparison with native insulin, putatively because of a slower release of monomeric insulin from the oligomeric state.

We next examined the cytotoxicity of resveratrol and EGCG in the presence of insulin. Resveratrol did not affect the viability of HepG2 cells even at 80μM concentration whereas EGCG reduced noticeably their viability at higher concentrations (Figure 3c). Resveratrol and EGCG in the presence of insulin consistently showed minimal cytotoxicity in comparison with fibrillated insulin in the live/dead assay (Figure 3d).

### Resveratrol stabilizes the population of a hexameric species of insulin molecule

The observation that oligomers formed upon resveratrol treatment preserve well the biological activity of insulin, instigated us to characterize structurally the dominant oligomeric species stabilized by resveratrol so that the mechanism of resveratrol-mediated stabilization of insulin can be discerned. 3D representation of the AFM images revealed that (Figure S4), resveratrol induced the formation of a small, almost uniform population of oligomeric structures of insulin (Figure S4b), in contrast to EGCG (Figure S4c) and curcumin (Figure S4d)-induced oligomers. The lower value of the average thickness of insulin amyloid fibril formed in the absence of polyphenolic compound (Figure S4a) is consistent with the identification of the presence of two monomers per unit length of narrow insulin fibril in a previous study [9].

Dynamic light scattering (DLS) measurements demonstrated that, while EGCG and curcumin induced the formation of oligomeric forms much larger in size, resveratrol stabilized smaller oligomers similar in size to the native insulin (Figure S2b). The chromatographic assessment identified hexamers as the primary form (obtained virtually as a single peak) of not only the native but also resveratrol-stabilized insulin (Figure S2c), suggesting that resveratrol stabilizes insulin predominantly in a hexameric form, closely similar to the native hexamer conformation.

To understand the mechanism underlying its anti-fibrillation effect, we set out to determine the 3D structure of the major oligomeric form of resveratrol-treated insulin using cryo-EM. Isolated single units of insulin oligomeric species with distinct morphology were observed evenly dispersed in cryo-EM micrographs (Figure 4a). Single particle 3D reconstruction technique was employed to generate the 3D structure of the resveratrol-stabilized insulin molecule. The reference free initial models, generated using EMAN2 [29] and Frealign [30] image processing software, had a globular shape which showed overall resemblance with the published crystal structures of insulin hexamer. The 2D class averages also showed spherical nature of the oligomer (Figure 4b). The final 3D cryo-EM map (∼14Å gold standard resolution) generated using SPIDER [31] (no symmetry applied) showed a globular structure with three lobes bulging out (Figure 4e). The 2D back projections (Figure 4c) of the final 3D map showed a clear similarity with the 2D class averages. 2D projections (Figure 4d) of the low pass filtered crystal structure of insulin hexamer stabilized by m-cresol (PDB code: 1EV6) also showed spherical appearance but the sizes appeared to be slightly smaller in some of the views as compared to the 2D class averages. Although the overall shape of the 3D density map resembles the crystal structure of the insulin hexamer, the map looks (Figure 4e) swollen as compared to the crystal structure (Figure 4f). Fitting of the crystal structure of insulin dimer into each lobe of the tri-lobed structure of the density map (Figure 4g) accommodated the atomic structures satisfactorily into the density map, indicating that resveratrol likely stabilizes insulin molecules primarily in a less compact hexameric form, compared to the more compact insulin hexamer stabilized by phenolic derivatives.

**Figure 4:**
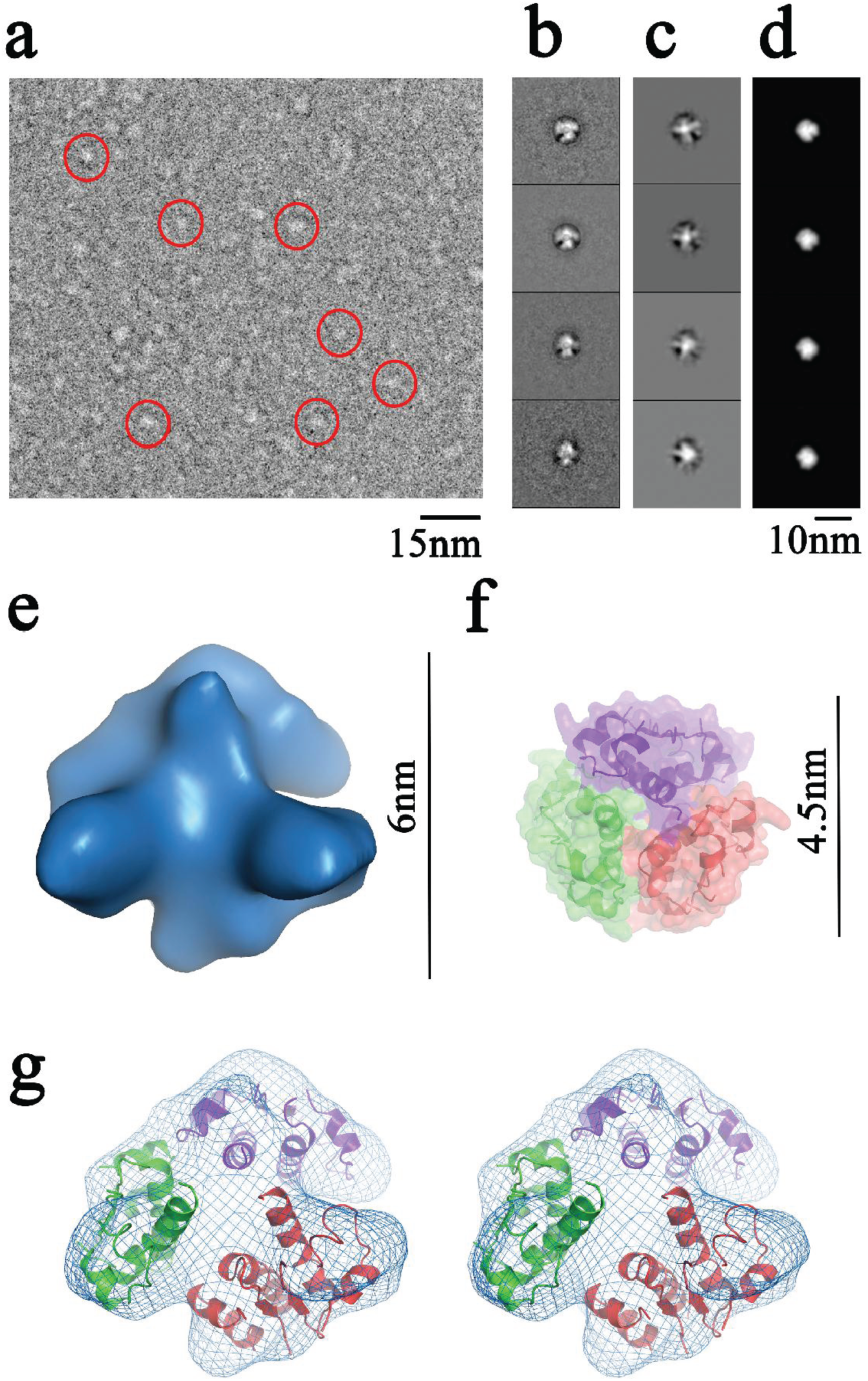
Cryo-EM single particle analysis of insulin-resveratrol complex. (a) A representative cryo-EM image of insulin-resveratrol complex showing distribution of the particles (marked with red circles). (b) 2D class averages of particle set processed in SPIDER. (c) Re-projections of the 3D structure in same Euler angles as determined for the selected 2D class averages. (d) Re-projection of the reference volume (converted from the crystal structure, PDB ID: 1EV6) in the same Euler angles. The projections of the reference volume appear to be slightly smaller. (e) Cryo-EM 3D density map of insulin in complex with resveratrol (generated in SPIDER) shows globular tri-lobed structure. (f) Crystal structure of the insulin hexamer containing phenolic derivatives (PDB: 1EV6) is shown in semitransparent surface view within which dimer structures are shown (three different colours) in ribbon representation. The crystal structure is more compact compared to the resveratrol-stabilized insulin. (g) Stereo view of the density map (blue mesh surface) where atomic structures of three dimers (ribbon diagrams in three different colours) are fitted into the lobes of the density map. Each lobe nicely accommodates a dimer of insulin molecule suggesting that resveratrol likely stabilizes insulin in a hexameric form.

Interestingly, when we performed refinement of each dimeric unit (without any ligand) of the hexamer crystal structure (using GalaxyWeb server), we found that the refined molecular model of the insulin hexamer showed an expanded conformation compared to the hexamer crystal structure (Figure S1f) indicating that an intrinsic tendency of relaxation is likely present in the hexamer structure of insulin (closely compact due to the presence of phenolic derivatives).

### Structural characterization of a resveratrol-stabilized oligomeric form of insulin

We performed an intrinsic fluorescence quenching experiment to determine interaction patterns of resveratrol with native insulin,. It was observed that intrinsic fluorescence level of insulin was decreased gradually in the presence of increasing concentration of resveratrol (Figure S2d).

Isothermal Titration Calorimetry (ITC) estimation further showed the specific nature of resveratrol binding to insulin. The stoichiometry for insulin titrated against resveratrol, suggested (Figure 5a) binding of three ligands per protein molecule. Since native insulin stays in hexameric form, as determined previously by chromatography (Figure S2c), it is conceivable that each hexamer binds to three resveratrol molecules. The thermodynamic parameters indicated the low binding affinity of resveratrol towards insulin (Kd in the micromolar range, ΔH: −5.45 kcal/mol) and further justified the easy release of active monomeric insulin.

**Figure 5:**
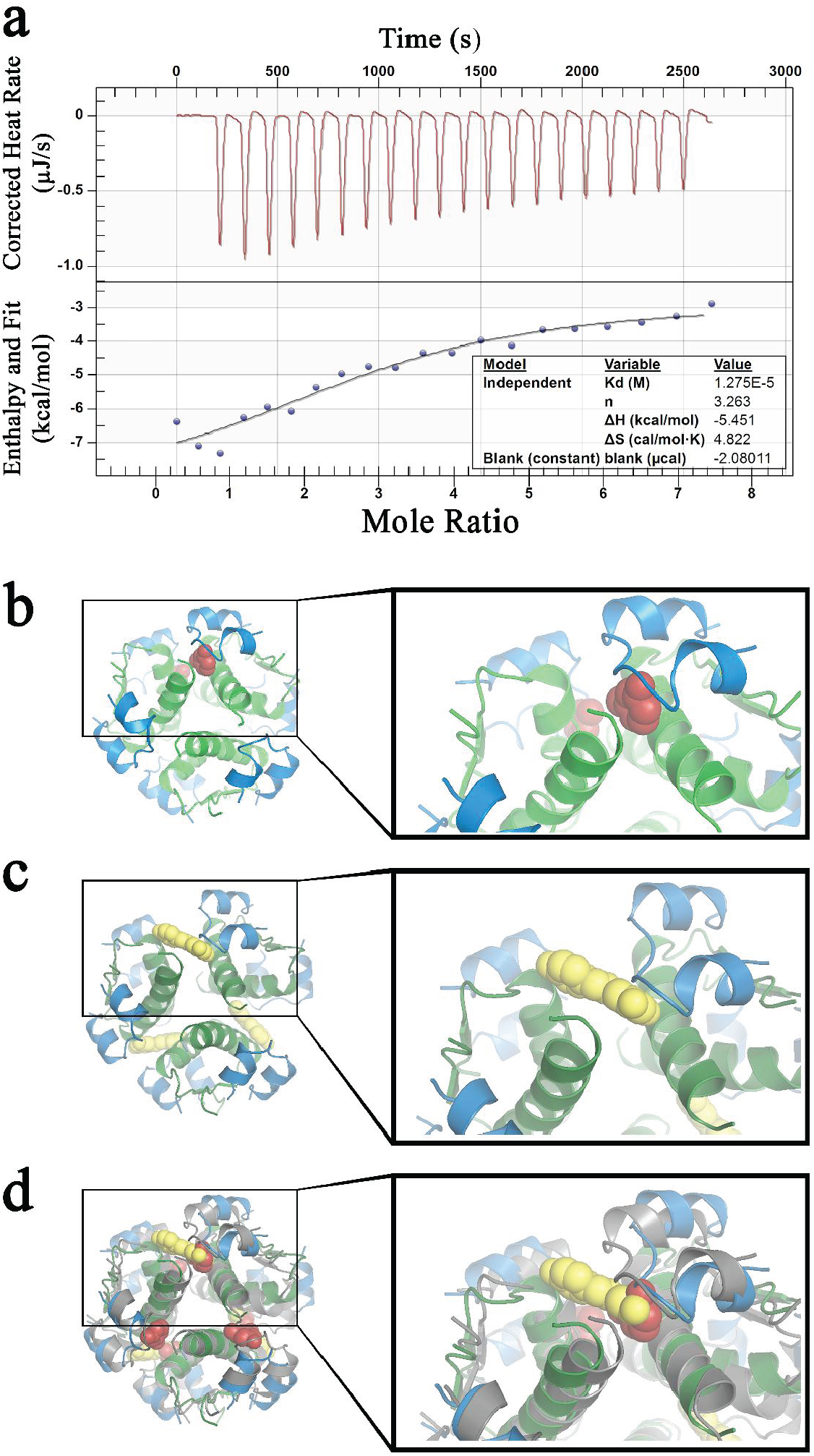
Analyses of ligand interactions with the insulin hexamer. (a) Interaction pattern of resveratrol with native insulin determined using Isothermal Titration Calorimetry (ITC). Insulin (20μM) is titrated against resveratrol (200μM). Equilibrium molar dissociation constant (Kd), insulin-resveratrol binding ratio (n), enthalpy (ΔH) and entropy changes (ΔS) are shown in the picture. (b) Binding sites of phenolic derivatives, m-cresol (shown in red) in the crystal structure of insulin hexamer (PDB: 1EV6). (c) Docked positions of resveratrol (shown in yellow) inside the crystal structure of insulin hexamer (following refinement in GalaxyWEB server) resulted from PatchDock. (d) Superposition of structures of (b) and (c) show overlapping positions of m-cresol and resveratrol. Right panels (in a, b, c) show the close up views of the protein-ligand interactions.

In order to decipher the binding sites of resveratrol within insulin hexamer, we performed docking studies on the atomic structure of insulin hexamer using commercially available docking software. We used the refined model structure (Figure S1f) of the insulin hexamer for docking in PatchDock server [32]. Top ten docked structures predicted by PatchDock showed that resveratrol molecules preferably occupy the interface of two dimeric modules (Figure 5). Crystallographic studies of insulin hexamer containing m-cresol molecules (or resorcinol, PDB code 1EVR) confirmed that m-cresol helps to stabilize the insulin hexamer structure in an active form [14, 33] and the crystal structures revealed that at least two phenolic derivatives (m-cresol/resorcinol) are placed at the interface of two dimers in a hexameric insulin molecule (Figure 5b), position-wise very similar to the resveratrol docking sites (Figure 5c). The strategic orientation of two phenolic groups in resveratrol molecule (Figure S1a) allows one resveratrol molecule to intercalate within a pair of insulin dimers to stabilize the hexameric structure, whereas two phenolic derivatives are required for such stabilization (Figure 5d). Notably, other docking servers e.g. SwissDock [34] and Autodock [35] also identified the pocket formed between two adjacent dimers in the hexameric structure as the potential binding region for resveratrol.

Combining the results of binding and docking studies we put forward a model of insulin hexamer stabilized by three resveratrol molecules, each accommodated at the interface of two adjacent dimers.

## Discussion

Injectable insulin remains the mainstay of diabetes therapy and developing newer formulations of insulin with better stability and stringently controlled duration of action are still the major challenges in clinics [36, 37]. Amyloidogenesis of insulin poses a stumbling block for reduced bioavailability during long-term storage [13]. Thus, identification of a nontoxic stabilizer that not only inhibits insulin fibrillation but also preserves its bioactivity has immense clinical importance.

Here we found that resveratrol, a natural polyphenol, not only blocks the amyloid formation of insulin, but that the resveratrol-bound insulin retains biological activity. Bioactivity of resveratrol-stabilized insulin both *in vitro* and *in vivo* clearly suggests enriched bioavailability of the active insulin monomer from resveratrol-stabilized insulin in comparison with native insulin undergoing fibrillation. Together, these findings reveal resveratrol as a potent insulin stabilizer showing anti-aggregating property by stacking insulin monomers into a stable hexameric form that fully retains its physiological activity.

Pharmaceutically formulated insulin contains phenolic compounds in combination with zinc ion (Zn^2+^) to stabilize insulin in hexameric form [14]. Structural studies manifested that some phenolic additives such as phenol, *m*-cresol or resorcinol promote subtle change in conformation of 1-8 residues of the shorter chain (chain B) from extended to helical structure and induce the transition of a hexameric assembly of insulin from inactive, less stable tensed (T) (T_6_ hexamers) state, to more active and stable relaxed (R) conformational state (R_6_ insulin) [14, 15, 38-40]. However, phenolic excipients are toxic in nature [16, 17]. This concern prompted us to think about resveratrol as a potent nontoxic insulin stabilizing agent and a prospective substitution for *m*-cresol for stabilizing insulin in a bioactive form.

Corollary investigation using structural characterization by Cryo-EM depicted a possible mechanism of resveratrol-mediated anti-fibrillation through stabilizing an active hexameric form of insulin, similar to the native insulin hexamer conformation, in a fibrillation-inducing condition. Analysis of the crystal structures revealed that insulin dimers are assembled and stabilized in a hexameric form in the presence of phenolic compounds and Zn^2+^ ion. Phenolic derivatives occupy the interface of adjacent dimers. In other words, insulin hexamer may be described as a trimer of dimers. Resveratrol apparently occupies the same position as the phenolic derivatives. However, in comparison to phenolic derivatives, resveratrol packs insulin dimers loosely into a trimeric form. An in-depth analysis by high-resolution structural characterization of resveratrol-stabilized insulin is imperative to delve deep into the mechanistic basis of insulin stabilization, which in turn will enhance the scope of new therapeutic formulation for insulin.

## Methods

### Preparation of insulin fibrils

Recombinant human insulin (hINS) powder (Sigma Aldrich) was dissolved in 20 mM HCl (pH 2), and the pH of the solution was immediately adjusted to 8 using 100 mM NaOH. Before incubation, the insulin solution was diluted with PBS (137 mM NaCl, 2.7 mM KCl, 8.1 mM Na_2_HPO_4_ and 1.47 mM KH_2_PO_4_, pH 7.4) at a concentration of 100 μM in the absence and in the presence of equimolar, sub-stoichiometric (25 μM and 50 μM), and super-stoichiometric (200 μM) concentration of polyphenols resveratrol (RES), Curcumin (CUR) and Epigallocatechin gallate (EGCG). Unless otherwise noted, fibrillation was induced by shaking the solution at 300 rpm, at 65°C for 24 h. All the buffer and insulin solutions were sterilized with a 0.22-μm filter (Millipore). Insulin concentration was measured by UV absorption at 276 nm by using an extinction coefficient of 1.0675 cm^−1^ (mg/ml) ^−1^.

### Thioflavin T assay

Thioflavin T (Th-T) dye binds specifically to the cross-β sheet structure of amyloid fibers and fluoresces more intensely in bound state [7]. After 24 h of incubation at 65°C in PBS (pH7.4), all samples (with or without polyphenols) were kept at 37°C for up to 14 days and ThT fluorescence values were measured at different days as indicated in the Figure 2. The molar concentration range of polyphenols RES, CUR and EGCG used in the insulin (100 μM) fibrillation experiment was 25 μM to 200 μM. Th-T was dissolved in mili-Q water, and its concentration was determined by UV absorbance at 412 nm using molar extinction coefficient of 36,000 M^−1^ cm^−1^. To obtain the fluorescence spectra, insulin fibrils were diluted to a final concentration of 0.5 μM in PBS containing 10 μM Th-T. Th-T fluorescence was measured in a quartz cuvette, using spectrofluorometer (F-7000) set at an excitation wavelength of 450 nm and emission wavelength set at 485 nm.

### Time-Course Experiments

100 μM of insulin in PBS solution was incubated with or without polyphenol at 65°C for different time intervals. Fibrillation was confirmed by Th-T assay and the amount of unreacted insulin was measured for each of the 24 h incubated samples. To measure the concentration of unreacted insulin, each sample was centrifuged at 16,000 g for 45 min, and soluble insulin concentration in the supernatant was determined by the Bradford protein assay at 595 nm.

### Fluorescence Measurement

Insulin contains Tyrosine residues (two in each of the chains A and B). Intrinsic fluorescence spectra of insulin was monitored on a spectro fluorimeter (Hitachi, F-7000) equipped with a circulating bath. Native insulin of 30 μM in 1 cm quartz cuvette was titrated with the successive additions of 10 mM resveratrol. Fluorescence emission spectra of insulin were measured at 310 nm with the excitation at 280 nm at 298K in the absence and presence of 2, 4, 8,16,30,60,100 and 120 μM of resveratrol.

### Isothermal titration calorimetry (ITC)

The interaction studies of native insulin with resveratrol were performed on Affinity ITC (TA Instruments, USA). Titration was performed in presence of 20 μM native insulin and 200 μM of resveratrol in PBS (pH 7.4) at 298K. All the solutions were extensively degassed before measurement to avoid bubble formation inside the colorimeter cell. During titration 2 μl of the resveratrol was injected to insulin containing cell at an interval of 300s. The buffer-resveratrol scan was subtracted from the insulin-resveratrol titration to correct the heat effects. The data were analysed and fitted by Nano Analyze software. Thermodynamic parameters were extracted for insulin-resveratrol interaction using the same software.

### Atomic Force Microscopy (AFM)

Insulin samples (100 μM) in the absence and in the presence of 100 μM resveratrol were incubated in PBS (pH 7.4) at 65°C for 24 h and then diluted 25 times. A small aliquot (5 µl) of the sample was collected and placed onto an even layered mica sheet for 10 minutes. The mica sheet was then washed with mili-Q pure water three times and air dried. AFM images were acquired in the air in different regions of each sample and representative images are reported. All experiments were performed in at least triplicates to validate the accuracy of the results.

### Transmission Electron Microscopy (TEM)

100 μM of insulin samples were subjected to fibrillation in the absence and presence of three different polyphenols independently. After 24 h of incubation, for TEM, 5 μl aliquots of the samples were applied onto the copper grid coated with a carbon film (300 meshes) and one drop of 2% uranyl acetate was placed on the grid. The excess water was removed carefully with filter paper, the grid was left to air dry and the images were taken with TECHNAI T12 TEM instrument.

### Confocal Microscopy

Samples were prepared by incubating 100 μM insulin in PBS (pH 7.4), with or without equimolar polyphenols (RES, EGCG, and CUR), at 65°C for 24 h. Aliquots (5µl) of each samples were mixed with 5μl PBS containing 10 μM Th-T and placed on clean glass slides and dried inside laminar airflow at room temperature in the dark. Th-T fluorescence of insulin samples was observed in a confocal microscope (OLYMPUS FV10i) using 405 nm band pass filter.

### Dynamic Light Scattering (DLS)

DLS was carried out at 25°C with a Malvern Zetasizer instrument equipped with temperature control. The measurement duration was 15 s, and 11 measurements were averaged for each analysis. Samples were prepared by incubating 100 μM insulin in PBS (pH 7.4) with equimolar polyphenols (RES, EGCG, and CUR) at 65°C for 24 h. After 24 h of incubation sample was analyzed by DLS immediately. The hydrodynamic size of insulin samples were acquired and plotted against percent mean volume. To ensure repeatability and statistics, each sample was measured in triplicate.

### Size Exclusion Chromatography (SEC)

SEC was performed using a Protein-pak 125 column from Waters. Insulin (100 μM) was incubated in PBS (pH 7.4) with equimolar polyphenols (RES and EGCG) at 65°C for 24 h. The SEC column was equilibrated with 10 mM sodium phosphate buffer containing 150 nm Nacl. Aliquots of 50 μl of each sample were loaded on the column and monitored at 280 nm. Bovine serum albumin (66.5 kD), ovalbumin (42.7kD), insulin hexamer (34.84 kD) and insulin monomer (5.8 kD) were run as markers prior to the loading of samples.

### Cryo-electron microscopy and image processing

Cryo-grids were prepared using Vitrobot Mark IV (FEI, USA). The samples (5μl) were deposited onto the 300 mesh Quantifoil R2/2 holey copper grids (washed and glow discharged prior to sample deposition), blotted (with constant temperature and humidity during the process of blotting), plunge frozen in liquid ethane and subsequently stored in liquid nitrogen. Images were recorded under low-dose conditions in Tecnai G2 Polara FEG-electron microscope (FEI) operating at 300 kV. Images were collected with 4K × 4K ‘Eagle’ charge-coupled device (CCD) camera (FEI, USA) at 78894 X magnification, resulting in a pixel size 1.89 Å at the specimen level.

Particle images were manually boxed with the EMAN2[41] program e2boxer.py. Defocus of the particles in each frame average was automatically determined by EMAN2 program e2ctf.py. Particles were then phase flipped and formed stack files for further processing. Two-dimensional (2D) reference-free class averages were computed by e2refine2d.py on the particle image sets. Same set of raw particles were subjected to classification independently in cisTEM (Frealign) [30, 42], and Xmipp [43] (implemented in SCIPION cryo-EM image processing framework (http://scipion.cnb.csic.es/m/home/). About 11000 particles were used in the particle dataset. Initial models were created in EMAN2 and Frealign which look similar to the crystal structure of the insulin hexamer.

We have generated final 3D density map (∼14Å resolution, Figure S5) in SPIDER (using new version where ‘gold standard’ refinement protocol has been implemented) using low pass filtered crystal structure (filtered to ∼25Å) as the initial model. To further validate the 3D structure of the oligomer, the 3D density map was also generated in Frealign (available in SCIPION) and EMAN2 using same particle set, which also resulted very similar final maps in shape and sizes.

### Modelling and docking

Fitting of the insulin dimer structure in each of the lobes of cryo-EM density map was done initially in Pymol (Delano Scientific) manually and then the model was subjected to molecular dynamics based fitting in MDFF[44] treating each of the dimer as a rigid body. ‘Refine’ module in GalaxyWeb server [45] (http://galaxy.seoklab.org/) was used for refining the dimeric units of insulin hexamer crystal structure (PDB ID:1EV6). Removal of ligands (e.g. Zn^2+^ ion, m-cresol etc.) from the PDB structure was done prior to uploading it to the server. Refined model structure of the insulin hexamer and resveratrol 3D structure (taken from a crystal structure, PDB ID: 2JIZ) were used for docking in PatchDock server [32] which is a molecular docking algorithm based on shape complementarity principle. Docking in PatchDock using insulin crystal structure and resveratrol was also done which resulted in same docking positions of the ligand inside the hexameric structure of the protein. The docking performed using Autodock [46] and SwissDock servers [34] also identified the region at the interface of two dimeric units as the primary binding sites of resveratrol. For creating illustrations, Chimera [47] and Pymol were used.

### Cell lysis and Immunoblotting

Cells were rinsed with ice-cold PBS and total cellular protein was prepared with lysis buffer containing 50 mM Tris-HCl (pH 7.4), 100 mM NaCl, 1 mM EDTA, 1 mM EGTA and 1% Triton X-100 with protease and phosphatase inhibitor cocktails both from (Millipore, Billierica, MA, USA). The soluble fractions of cell lysates were isolated by centrifugation at 15000g for 15 minutes at 4^°^C. Protein samples (40 μg/lane) were resolved by 10% SDS-PAGE and transferred to Immobilon-P membranes (Millipore, Billierica, MA, USA) by wet transfer (Trans-Blot, Bio-Rad) at 90 V for 3 h. For immunodetection, membrane was blocked with 5% non-fat milk in PBS with 0.1% Tween 20 for 1 h, followed by incubation with specific primary antibody at 4^°^C overnight and with horseradish peroxidase-labeled secondary antibodies for 1 h at room temperature. Signals were detected by chemiluminescence using Luminata Classico Western HRP substrate (Millipore, Billierica, MA, USA) and scanned using a ChemiDocMP System (Bio-Rad Laboratories, Hercules, CA). Detailed list of antibodies is provided below.

### Animal Study

All experiments with animals were performed under the approved Institutional Animal Ethics Committee (Approved by CPCSEA, India) protocol. Eight to ten week old, male wild-type BALB/c mice were kept at ambient temperature (22 ± 1°C) with 12:12 h light/dark cycles and fed standard chow diet (4.3% lipid and 70% carbohydrate). Mice were fasted for 6 hours and fasting blood glucose was measured using a calibrated glucometer by taking one drop of blood from the tail tip cut. Intraperitoneal injections of insulin (0.5 U/kg of body weight) were administered with a 27 G needle. Blood samples were taken from the initial tail cut at 30, 60 and 120 minutes after intraperitoneal insulin injection. This experiment was carried out for four different groups: insulin, fibrillated insulin, fibrillated insulin+EGCG and fibrillated insulin+RES. Insulin Tolerance Test has been replicated in five mice per group.

### Live Dead Assay

HepG2 cells were harvested in DMEM and 0.5 ml of 0.4% Trypan Blue solution (w/v) was added to the suspension of cells. A small amount of Trypan Blue-cell suspension mixture was placed on the hemocytometer for counting. Cells in all the chambers were counted and five replicate readings were taken. Approximately 2.5 × 10^5^ cells were counted and plated into each well to determine cell viability. Next day, cells were treated with 40 uM of either PBS, fibrillated insulin, fibrillated insulin with RES or fibrilated insulin with EGCG. After 24 hrs of treatment, cell viability was assessed for respective treatments using the Live/Dead viability kit (Invitrogen, Renfrew).

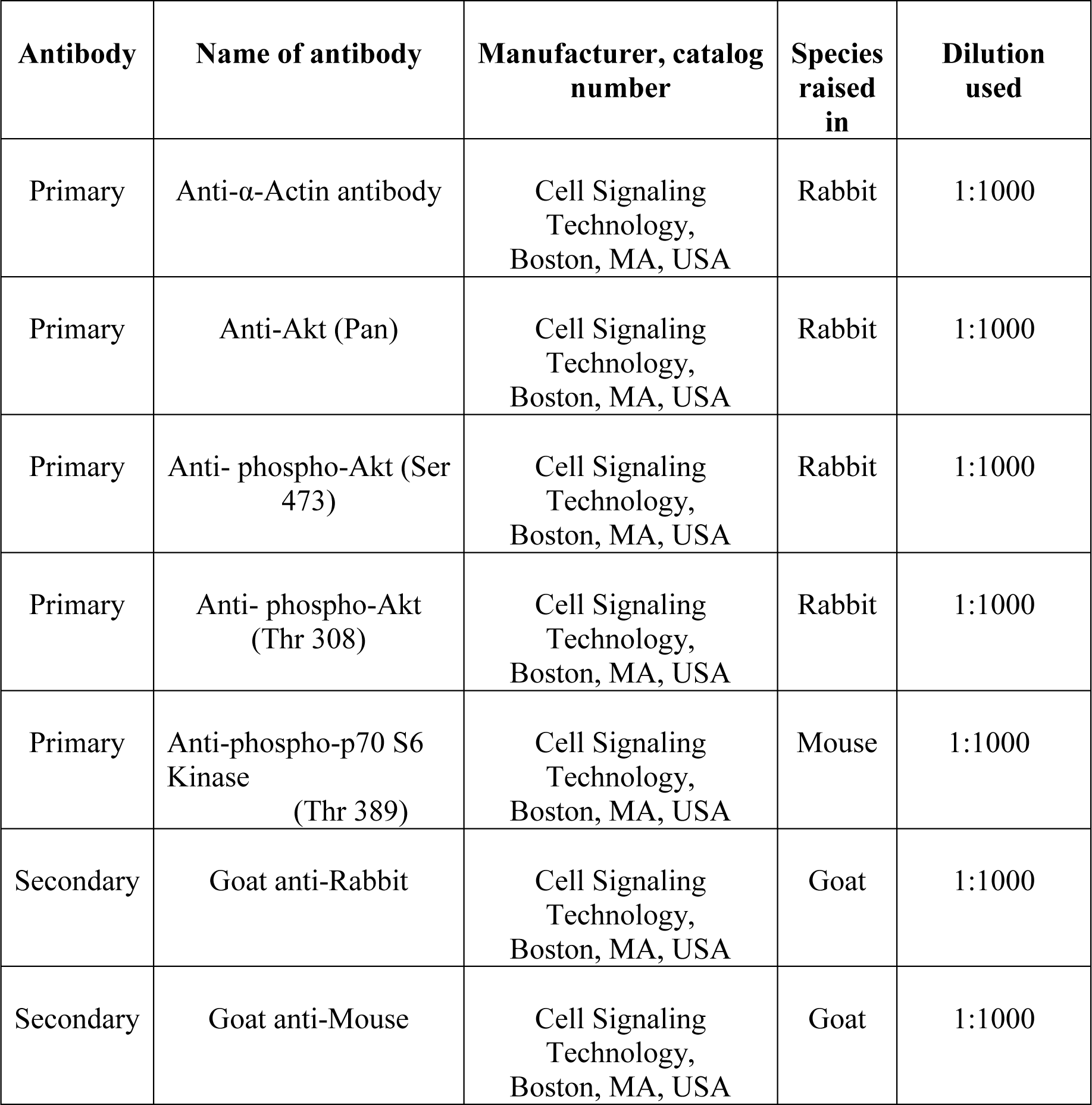

### Cytotoxicity assay

HepG2 cells grown in DMEM medium were harvested and 0.4% Trypan Blue solution was added to the cell suspension for counting using hemocytometer. Approximately10^5^ cells/100 μl of medium per well were plated in 96 well polystyrene plates. Cells were treated with resveratrol and EGCG at concentrations of 20, 40 or 60 uM for 24 hrs. After 24 hrs of treatment, cell viability was determined with the addition of 10 μl of 5 mg/ml MTT to each well. After incubation for 4 hrs at 37°C, the medium was aspirated from each well and 100 μl of DMSO was added per well. Plates were agitated at 25°C for 10 min and absorbance was recorded at 590 nm using a multi-well plate reader (Biotek-Epoch; Biotech Instruments). The average absorbance value of five replicate wells was used for each set and each experiment was repeated thrice. In this assay, cells treated with PBS and no test samples but MTT served as positive control. Cells without MTT served as blank.

### Statistics

Statistical values are presented as the mean ± SD. A two-tailed Student’s t-test was used to calculate the p-values.

## Acknowledgements

This work was supported by SERB, DST (India) sponsored project, DBT contingency fund for Research Associate (DBT-RA, India), and CSIR-Indian Institute of Chemical Biology, Kolkata, India. BP sincerely thanks DBT for providing Research associate Fellowship. We acknowledge the Central Instrument Facility (CIF) (and all the technical staff associated with it) of CSIR-IICB. We thank Mr. Chiranjit Biswas for cryo-EM data collection. DD and SB acknowledge UGC and CSIR (India), respectively, for Senior Research Fellowship.

## Author contributions

BP and JS conceived the project and designed the experiments. BP carried out biochemical, biophysical experiments and 3D image processing of cryo-EM data. DD performed cell-based and animal experiments under the supervision of PC. SB assisted in cryo-EM data processing. BP, DD, PC and JS analyzed the data and wrote the paper.

*There are no conflicts of interest to declare

## Supplementary Figures

**Figure S1:**
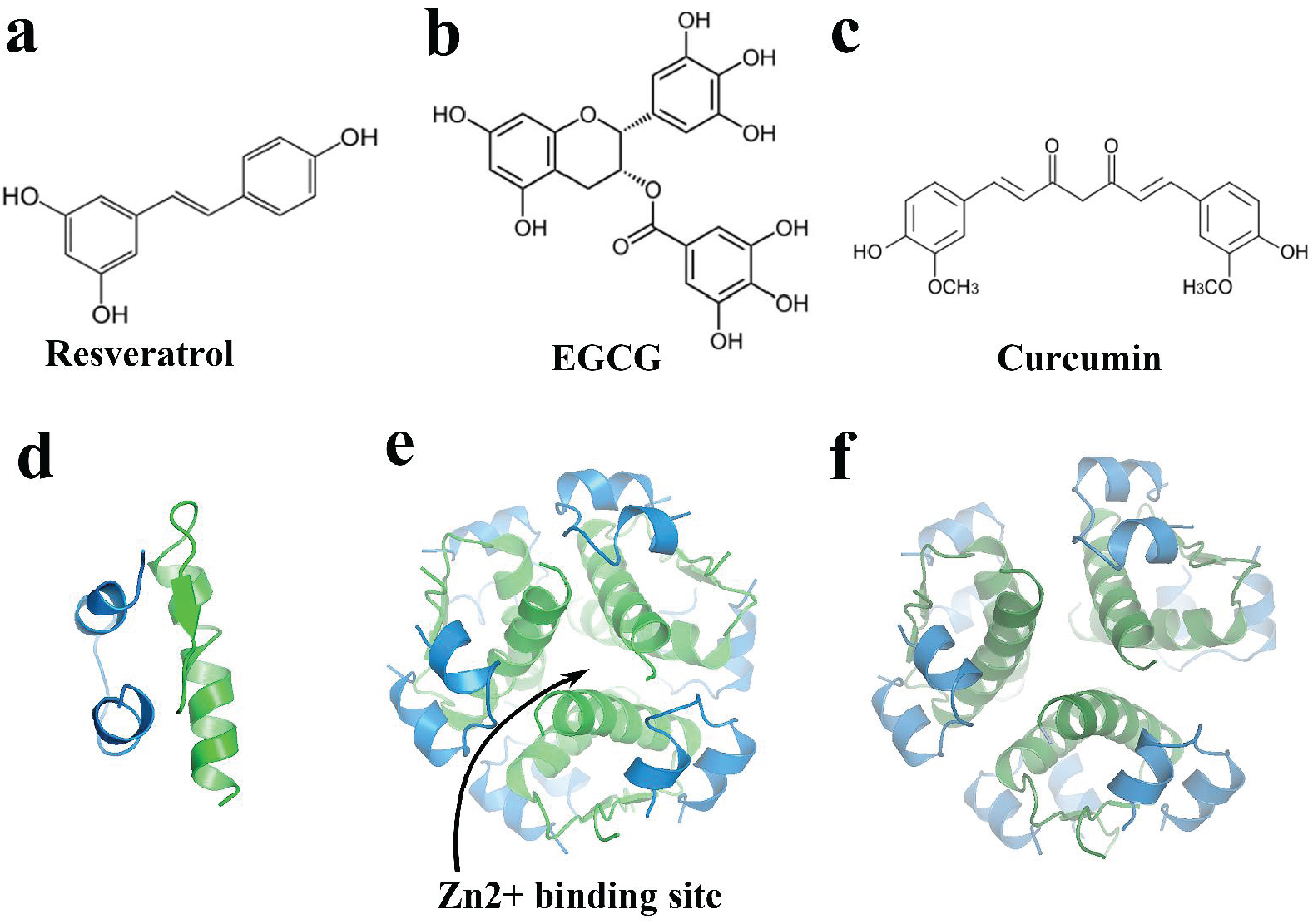
Structures of polyphenols and insulin in different forms. Chemical structures of (a) resveratrol, (b) EGCG, and (c) curcumin. (d) The native monomeric insulin has two chain, short (blue) and long (green), linked to each other by two disulfide bonds. (e) Insulin hexamer and Zn^2+^ binding site within it. (f) Insulin hexamer structure without any small molecules) when refined in GalaxyWEB shows slight relaxation in the packing of 3 dimers.

**Figure S2:**
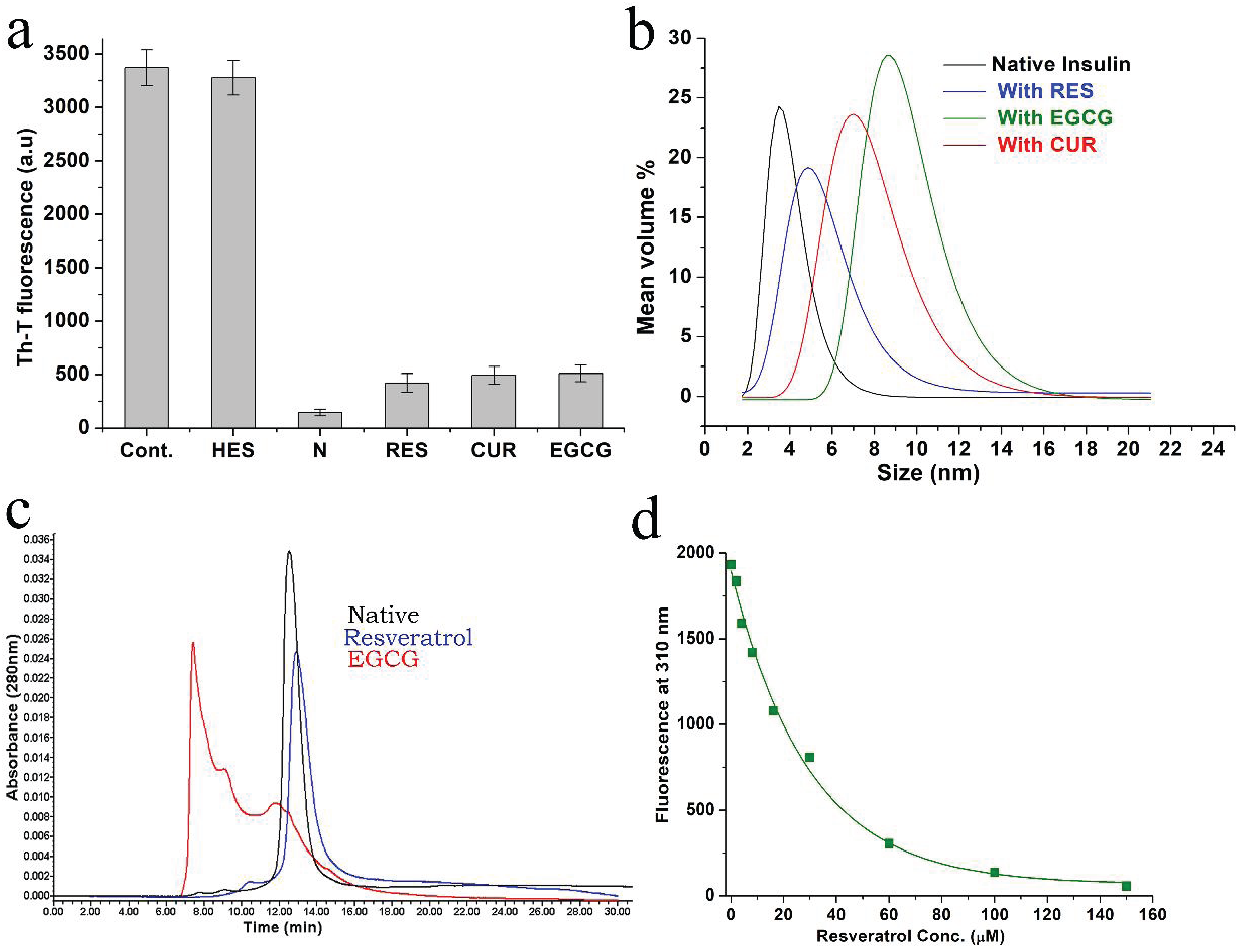
Effects of small molecules on insulin aggregation. (a) Th-T fluorescence of insulin fibrillation after 24 hrs of incubation at 65^0^C in PBS (pH7.4) in the absence and presence of flavanone glycoside-hesperidin (HES) and three native polyphenols, resveratrol (RES), curcumin (CUR) and epigallocatechin gallate (EGCG). Native (N) insulin fluorescence was included in the bar diagram as standard Th-T fluorescence spectra. (b) Dynamic light scattering (DLS) data shows the mean volume percent respect to size (nm) of the native and polyphenols treated insulin as indicated by different color codes in the picture. In the experiment, polyphenols treated samples were prepared by incubation of insulin (100μM) with polyphenols (100μM) for 24h. (c) Elution profile of native insulin (Native), resveratrol and EGCG treated insulin in Size exclusion chromatography has been compared. Proteins standard BSA (MW 66.5 kDa), ovalbumin (MW 42.7kDa), insulin hexamer (MW 34.84 kDa), trypsin (MW 23.3 kDa) and insulin monomer (MW 5.8 kDa) were loaded on HPLC and their molecular weight vs retention time inside the chromatography column is plotted to generate the standard curve. The retention time of native insulin (Native) in PBS and resveratrol-treated insulin (resveratrol) are same as shown in the chromatogram. The molecular weight of native and resveratrol-treated insulin is 32.76 kDa and 31.5 kDa, respectively when calculated from the standard graph (using Y= −0.0967X +2.7066 equation, X= retention time of unknown). However, EGCG treated insulin eluted before the native insulin or resveratrol-treated insulin. (d) Insulin short and long chains contain Tyr residues (two Tyr/chain). Quenching of intrinsic fluorescence of insulin when titrated with 10 mM resveratrol is shown.

**Figure S3:**
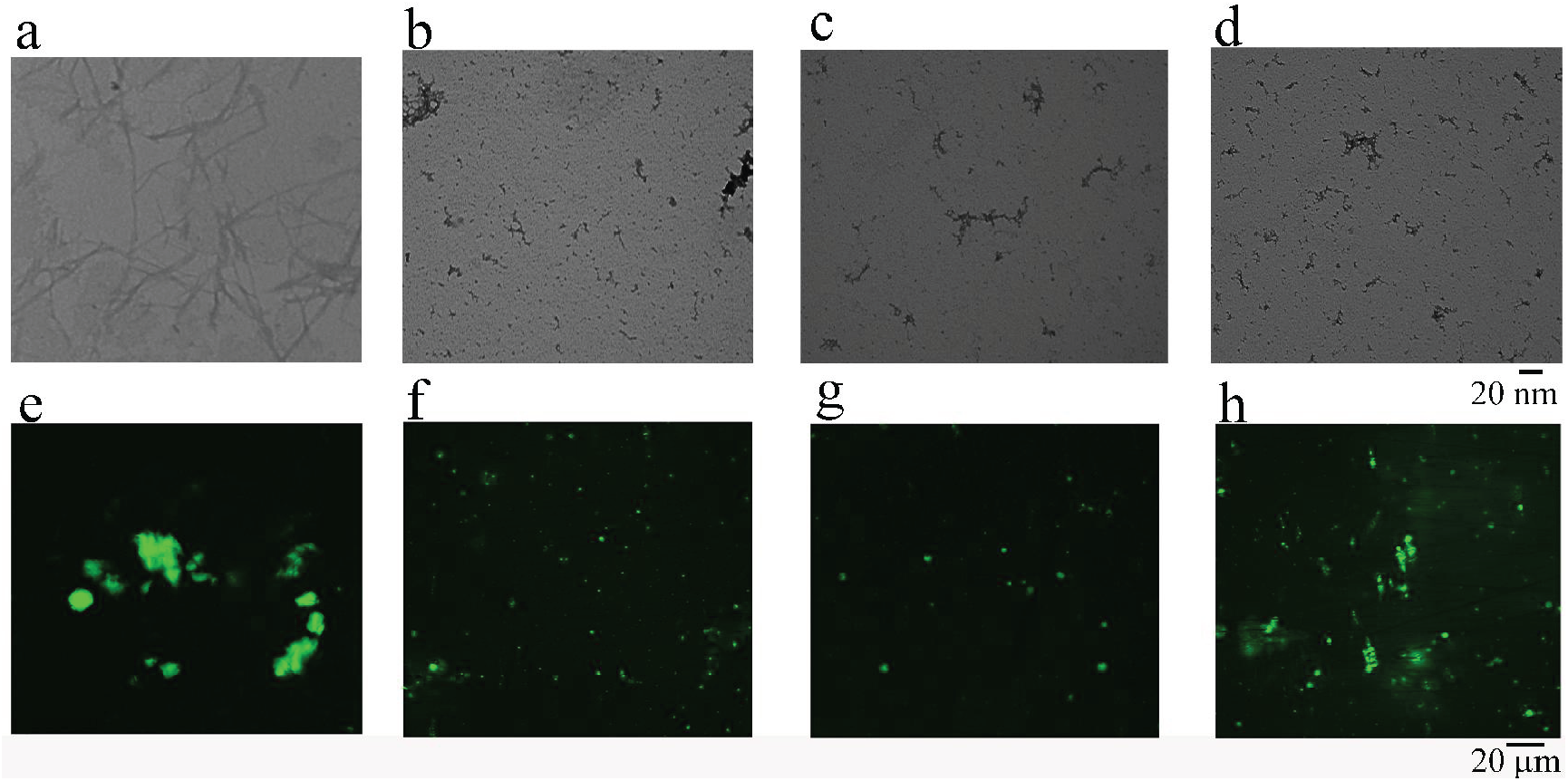
Dose-dependent responses of resveratrol on insulin aggregation. TEM images of insulin (100 μM) fibrillation in the absence (a) and presence of (b) 200 μM (1:2), (c) 50 μM (1:0.5), and (d) 25 μM (1:0.25) resveratrol are shown. Confocal images of insulin fibrillation shows in the absence (e) and presence of 200 μM (f), 50 μM (g) and 25 μM (h) resveratrol. These results suggest that the dose for effective anti-aggregation property of resveratrol, is when insulin:resveratrol ration is 1:0.5 or higher.

**Figure S4:**
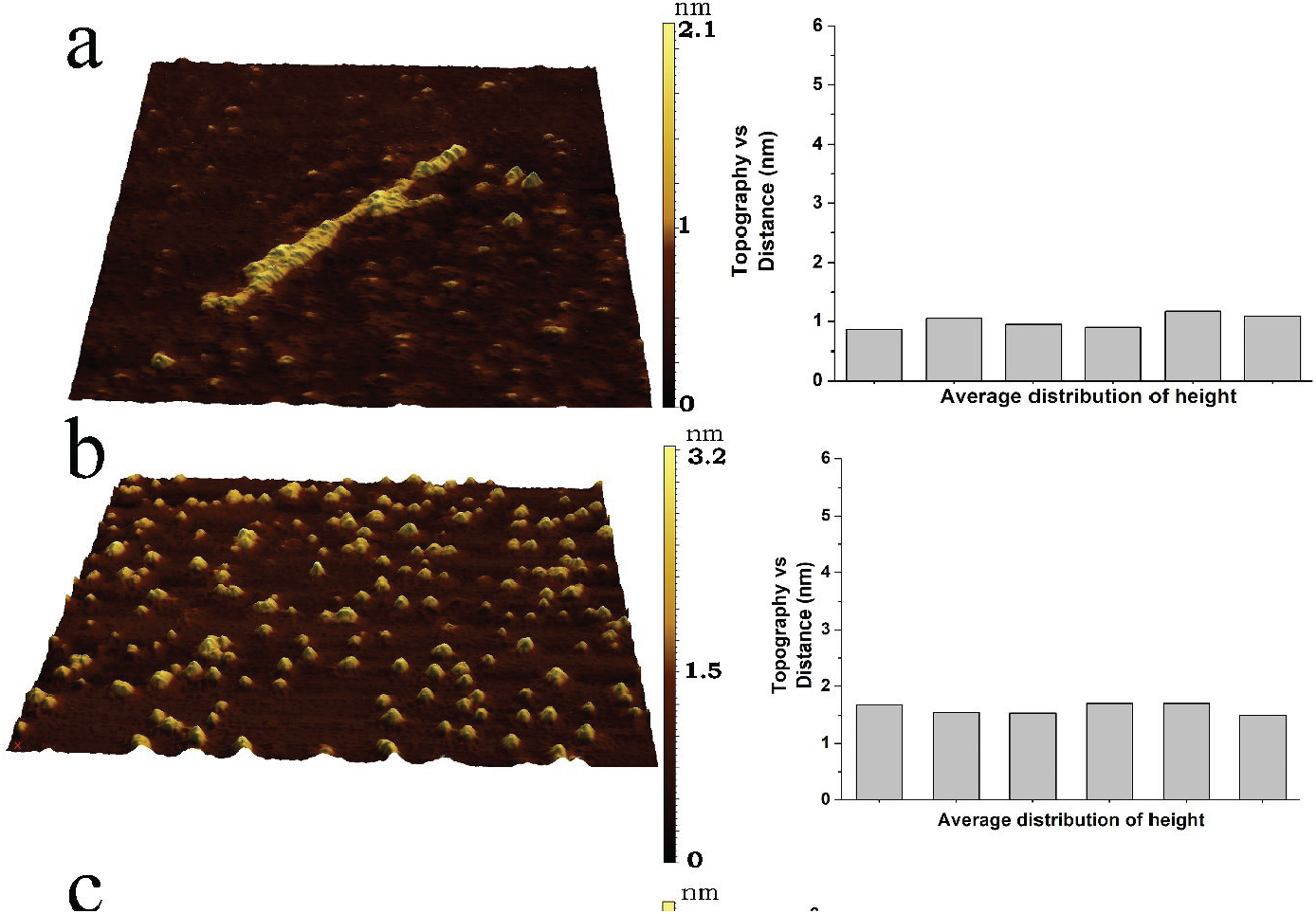
Representation of the AFM images in 3D. 3D view of the AFM images of insulin without any polyphenol (a), with RES (b), EGCG (c), CUR (d). Right panels show the mean distribution of molecular heights of the amyloid fibrils formed without polyphenols, and insulin oligomers stabilized by the polyphenols.

**Figure S5:**
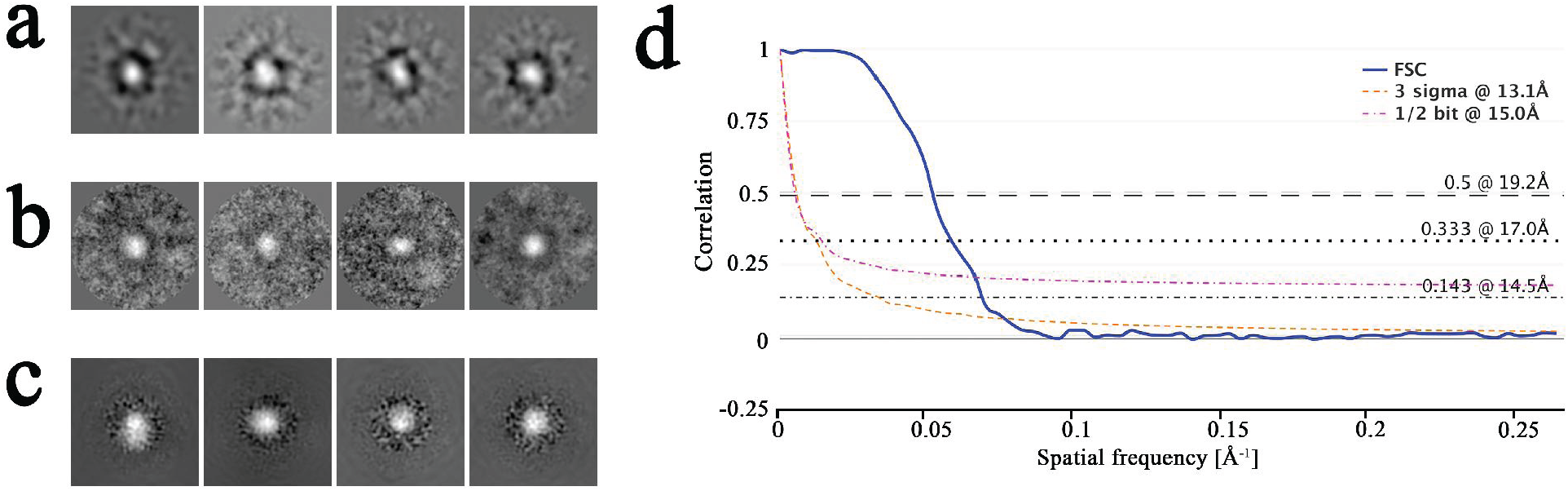
Validation of 3D cryo-EM structure of the insulin-resveratrol complex. (a) 2D class averages of the particles set generated by EMAN2 (a), Xmipp (b), and cisTEM (Frealign) (c) show a very similar spherical feature. (d) Fourier shell correlation (FSC) plot (generated in EMDB web server (https://www.ebi.ac.uk/pdbe/emdb/validation/fsc/)) of the single particle 3D reconstruction (done using newly developed gold standard protocol in SPIDER) of resveratrol-stabilized insulin.

## References

1. Brange, J. & Langkjoer, L. (1993) Insulin structure and stability, Pharm Biotechnol. 5, 315–50.

2. Blundell, T. L., Cutfield, J. F., Cutfield, S. M., Dodson, E. J., Dodson, G. G., Hodgkin, D. C., Mercola, D. A. & Vijayan, M. (1971) Atomic positions in rhombohedral 2-zinc insulin crystals, Nature. 231, 506–11.

3. Bryant, C., Spencer, D. B., Miller, A., Bakaysa, D. L., McCune, K. S., Maple, S. R., Pekar, A. H. & Brems, D. N. (1993) Acid stabilization of insulin, Biochemistry. 32, 8075–82.

4. Dunn, M. F. (2005) Zinc-ligand interactions modulate assembly and stability of the insulin hexamer -- a review, Biometals. 18, 295–303.

5. Dodson, G. & Steiner, D. (1998) The role of assembly in insulin’s biosynthesis, Curr Opin Struct Biol. 8, 189–94.

6. Xu, Y., Yan, Y., Seeman, D., Sun, L. & Dubin, P. L. (2012) Multimerization and aggregation of native-state insulin: effect of zinc, Langmuir. 28, 579–86.

7. Ratha, B. N., Ghosh, A., Brender, J. R., Gayen, N., Ilyas, H., Neeraja, C., Das, K. P., Mandal, A. K. & Bhunia, A. (2016) Inhibition of Insulin Amyloid Fibrillation by a Novel Amphipathic Heptapeptide: MECHANISTIC DETAILS STUDIED BY SPECTROSCOPY IN COMBINATION WITH MICROSCOPY, J Biol Chem. 291, 23545–23556.

8. Lee, C. C., Nayak, A., Sethuraman, A., Belfort, G. & McRae, G. J. (2007) A three-stage kinetic model of amyloid fibrillation, Biophys J. 92, 3448–58.

9. Ivanova, M. I., Sievers, S. A., Sawaya, M. R., Wall, J. S. & Eisenberg, D. (2009) Molecular basis for insulin fibril assembly, Proc Natl Acad Sci U S A. 106, 18990–5.

10. Horvath, I. & Wittung-Stafshede, P. (2016) Cross-talk between amyloidogenic proteins in type-2 diabetes and Parkinson’s disease, Proc Natl Acad Sci U S A. 113, 12473–12477.

11. Brange, J., Andersen, L., Laursen, E. D., Meyn, G. & Rasmussen, E. (1997) Toward understanding insulin fibrillation, J Pharm Sci. 86, 517–25.

12. Dische, F. E., Wernstedt, C., Westermark, G. T., Westermark, P., Pepys, M. B., Rennie, J. A., Gilbey, S. G. & Watkins, P. J. (1988) Insulin as an amyloid-fibril protein at sites of repeated insulin injections in a diabetic patient, Diabetologia. 31, 158–61.

13. Woods, R. J., Alarcon, J., McVey, E. & Pettis, R. J. (2012) Intrinsic fibrillation of fast-acting insulin analogs, J Diabetes Sci Technol. 6, 265–76.

14. Whittingham, J. L., Edwards, D. J., Antson, A. A., Clarkson, J. M. & Dodson, G. G. (1998) Interactions of phenol and m-cresol in the insulin hexamer, and their effect on the association properties of B28 pro --> Asp insulin analogues, Biochemistry. 37, 11516–23.

15. Derewenda, U., Derewenda, Z., Dodson, E. J., Dodson, G. G., Reynolds, C. D., Smith, G. D., Sparks, C. & Swenson, D. (1989) Phenol stabilizes more helix in a new symmetrical zinc insulin hexamer, Nature. 338, 594–6.

16. Jarosz-Chobot, P., Nowakowska, M. & Polanska, J. (2007) Seeking the factors predisposing to local skin inflammatory state development in children with type 1 diabetes (T1DM) treated with continuous subcutaneous insulin infusion (CSII), Experimental and clinical endocrinology & diabetes: official journal, German Society of Endocrinology [and] German Diabetes Association. 115, 179–81.

17. Weber, C., Kammerer, D., Streit, B. & Licht, A. H. (2015) Phenolic excipients of insulin formulations induce cell death, pro-inflammatory signaling and MCP-1 release, Toxicology reports. 2, 194–202.

18. Johansson, U. B., Adamson, U., Lins, P. E. & Wredling, R. (2005) Patient management of long-term continuous subcutaneous insulin infusion, Journal of advanced nursing. 51, 112–8.

19. Sanders, J. M., Bucher, J. R., Peckham, J. C., Kissling, G. E., Hejtmancik, M. R. & Chhabra, R. S. (2009) Carcinogenesis studies of cresols in rats and mice, Toxicology. 257, 33–9.

20. Boross, G. N., Shimura, S., Besenius, M., Tennagels, N., Rossen, K., Wagner, M. & Bode, J. W. (2018) Facile folding of insulin variants bearing a prosthetic C-peptide prepared by alpha-ketoacid-hydroxylamine (KAHA) ligation, Chemical science. 9, 8388–8395.

21. Stirpe, A., Pantusa, M., Rizzuti, B., De Santo, M. P., Sportelli, L., Bartucci, R. & Guzzi, R. (2016) Resveratrol induces thermal stabilization of human serum albumin and modulates the early aggregation stage, Int J Biol Macromol. 92, 1049–1056.

22. Gong, H., He, Z., Peng, A., Zhang, X., Cheng, B., Sun, Y., Zheng, L. & Huang, K. (2014) Effects of several quinones on insulin aggregation, Sci Rep. 4, 5648.

23. Mishra, R., Sellin, D., Radovan, D., Gohlke, A. & Winter, R. (2009) Inhibiting islet amyloid polypeptide fibril formation by the red wine compound resveratrol, Chembiochem. 10, 445–9.

24. Castro-Acosta, M. L., Stone, S. G., Mok, J. E., Mhajan, R. K., Fu, C. I., Lenihan-Geels, G. N., Corpe, C. P. & Hall, W. L. (2017) Apple and blackcurrant polyphenol-rich drinks decrease postprandial glucose, insulin and incretin response to a high-carbohydrate meal in healthy men and women, The Journal of nutritional biochemistry. 49, 53–62.

25. Costabile, G., Vitale, M., Luongo, D., Naviglio, D., Vetrani, C., Ciciola, P., Tura, A., Castello, F., Mena, P., Del Rio, D., Capaldo, B., Rivellese, A. A., Riccardi, G. & Giacco, R. (2018) Grape pomace polyphenols improve insulin response to a standard meal in healthy individuals: A pilot study, Clinical nutrition.

26. Yoshihara, H., Saito, J., Tanabe, A., Amada, T., Asakura, T., Kitagawa, K. & Asada, S. (2016) Characterization of Novel Insulin Fibrils That Show Strong Cytotoxicity Under Physiological pH, J Pharm Sci. 105, 1419–26.

27. Waugh, D. F. (1957) A mechanism for the formation of fibrils from protein molecules, J Cell Physiol Suppl. 49, 145–64.

28. Selivanova, O. M. & Galzitskaya, O. V. (2012) Structural polymorphism and possible pathways of amyloid fibril formation on the example of insulin protein, Biochemistry (Mosc). 77, 1237–47.

29. Tang, G., Peng, L., Baldwin, P. R., Mann, D. S., Jiang, W., Rees, I. & Ludtke, S. J. (2007) EMAN2: an extensible image processing suite for electron microscopy, J Struct Biol. 157, 38–46.

30. Grigorieff, N. (2007) FREALIGN: high-resolution refinement of single particle structures, Journal of structural biology. 157, 117–25.

31. Shaikh, T. R., Gao, H., Baxter, W. T., Asturias, F. J., Boisset, N., Leith, A. & Frank, J. (2008) SPIDER image processing for single-particle reconstruction of biological macromolecules from electron micrographs, Nature protocols. 3, 1941–74.

32. Schneidman-Duhovny, D., Inbar, Y., Nussinov, R. & Wolfson, H. J. (2005) PatchDock and SymmDock: servers for rigid and symmetric docking, Nucleic acids research. 33, W363–7.

33. Smith, G. D., Ciszak, E., Magrum, L. A., Pangborn, W. A. & Blessing, R. H. (2000) R6 hexameric insulin complexed with m-cresol or resorcinol, Acta Crystallogr D Biol Crystallogr. 56, 1541–8.

34. Grosdidier, A., Zoete, V. & Michielin, O. (2011) SwissDock, a protein-small molecule docking web service based on EADock DSS, Nucleic acids research. 39, W270–7.

35. Morris, G. M., Huey, R., Lindstrom, W., Sanner, M. F., Belew, R. K., Goodsell, D. S. & Olson, A. J. (2009) AutoDock4 and AutoDockTools4: Automated docking with selective receptor flexibility, Journal of computational chemistry. 30, 2785–91.

36. Jars, M. U., Hvass, A. & Waaben, D. (2002) Insulin aspart (AspB28 human insulin) derivatives formed in pharmaceutical solutions, Pharmaceutical research. 19, 621–8.

37. Teska, B. M., Alarcon, J., Pettis, R. J., Randolph, T. W. & Carpenter, J. F. (2014) Effects of phenol and meta-cresol depletion on insulin analog stability at physiological temperature, J Pharm Sci. 103, 2255–67.

38. Smith, G. D. & Dodson, G. G. (1992) Structure of a rhombohedral R6 insulin/phenol complex, Proteins. 14, 401–8.

39. Whittingham, J. L., Chaudhuri, S., Dodson, E. J., Moody, P. C. & Dodson, G. G. (1995) X-ray crystallographic studies on hexameric insulins in the presence of helix-stabilizing agents, thiocyanate, methylparaben, and phenol, Biochemistry. 34, 15553–63.

40. Smith, G. D., Ciszak, E. & Pangborn, W. (1996) A novel complex of a phenolic derivative with insulin: structural features related to the T-->R transition, Protein Sci. 5, 1502–11.

41. Tang, G., Peng, L., Baldwin, P.R., Mann, D.S., Jiang, W., Rees, I., Ludtke. S.J. (2007) EMAN2: an extensible image processing suite for electron microscopy J Struct Biol 157, 38–46.

42. Grant, T., Rohou, A. & Grigorieff, N. (2018) cisTEM, user-friendly software for single-particle image processing, eLife. 7.

43. Sorzano, C. O., Marabini, R., Velazquez-Muriel, J., Bilbao-Castro, J. R., Scheres, S. H., Carazo, J. M. & Pascual-Montano, A. (2004) XMIPP: a new generation of an open-source image processing package for electron microscopy, Journal of structural biology. 148, 194–204.

44. Trabuco, L. G., Villa, E., Schreiner, E., Harrison, C. B. & Schulten, K. (2009) Molecular dynamics flexible fitting: a practical guide to combine cryo-electron microscopy and X-ray crystallography, Methods. 49, 174–80.

45. Afgan, E., Baker, D., Batut, B., van den Beek, M., Bouvier, D., Cech, M., Chilton, J., Clements, D., Coraor, N., Gruning, B. A., Guerler, A., Hillman-Jackson, J., Hiltemann, S., Jalili, V., Rasche, H., Soranzo, N., Goecks, J., Taylor, J., Nekrutenko, A. & Blankenberg, D. (2018) The Galaxy platform for accessible, reproducible and collaborative biomedical analyses: 2018 update, Nucleic acids research. 46, W537–W544.

46. Morris, G. M., Huey, R., Lindstrom, W., Sanner, M. F., Belew, R. K., Goodsell, D. S. and Olson, A. J. (2009) Autodock4 and AutoDockTools4: automated docking with selective receptor flexiblity., J Computational Chemistry 16, 2785–91.

47. Pettersen, E. F., Goddard, T. D., Huang, C. C., Couch, G. S., Greenblatt, D. M., Meng, E. C. & Ferrin, T. E. (2004) UCSF Chimera--a visualization system for exploratory research and analysis, J Comput Chem. 25, 1605–12.

